# An Unsupervised Graph Embeddings Approach to Multiplex Immunofluorescence Image Exploration

**DOI:** 10.1101/2021.06.09.447654

**Authors:** Christopher Innocenti, Zhenning Zhang, Balaji Selvaraj, Isabelle Gaffney, Michalis Frangos, Jake Cohen-Setton, Laura A L Dillon, Michael J Surace, Carlos Pedrinaci, Jason Hipp, Khan Baykaner

## Abstract

Understanding the complex biology of the tumor microenvironment (TME) is necessary to understand the mechanisms of action of immuno-oncology therapies and to match the right therapies to the right patients. Multiplex immunofluorescence (mIF) is a useful technology that has tremendous potential to further our understanding of cancer patho-biology; however, tools that fully leverage the high dimensionality of this data are still in their infancy. We describe here a novel deep learning pipeline aimed to allow **Graph**-based **I**nspection of **T**issues via **E**mbeddings, **GraphITE**. GraphITE transforms mIF data into a graph representation, where unsupervised learning algorithms can be utilised to generate embeddings representing cellular ‘neighbourhoods’. The embeddings can be downprojected and explored for clustering analysis, and patterns can be mapped back to the image as well as interrogated for phenotypical, morphological, or structural distinctiveness. GraphITE supports the extraction of information not only on the phenotypes of individual cells or the relationships between specific cell types, but is able to characterize cell neighborhoods to look for more complex interactions, thereby allowing pathologists and data scientists to explore mIF data sets, uncovering patterns that are otherwise obscured by the high-dimensionality of the data. In this work, we showcase the current setup of the system, going from raw input data all the way to a user friendly exploration tool. Using this tool, we show how the data can be navigated in a way previously not possible.

## 1 INTRODUCTION

Immuno-oncology (IO) therapies have been shown to be highly effective in some cases, however patient response is known to be highly variable, with some patients failing to respond at all. Developing a deeper understanding of the tumor micro environment (TME) by characterizing phenotypes and cell locations is crucial for the development of anti-cancer therapeutic interventions. Multiplex immunofluorescence (mIF) imaging is increasingly being employed to gain insight into the TME, but interpretation of this data modality remains challenging. Both the power and the complexity of mIF imaging technology go hand-in-hand; it is the high-dimensionality that simultaneously captures spatial topology and rich phenotypic information that also makes data exploration difficult. To fully utilise the power of mIF, a method is needed to convert this complex, high-dimensional data into a lower-dimensional form wherein interesting patterns are made plain to pathologists.

Currently, pathologists use human cognition to identify unique patterns by toggling between biomarkers in an mIF image. Existing methods allow pathologists to view aggregated phenotypical information and cell interactions on a limited, generally one-to-one, basis. These methods have been largely descriptive and geared toward addressing particular hypotheses; to fully exploit mIF data a more holistic approach is required, one that leverages the spatial and phenotypic data prior to hypothesis-testing.

In the field of artificial intelligence (AI) and deep learning there exist such methods for converting complex, high-dimensional data into alternative, lower-dimensional representations that emphasise the otherwise hard to discern patterns spread across dimensions. A suite of tools known as unsupervised learning algorithms are appropriate for solving such problems; these are algorithms that aim to capture the structure of data, without reference to labels or outcomes.

In this work we propose a novel pipeline for converting individual cells (or whole tissues) from mIF images into graph (or cell level) embeddings. First a graph is constructed to represent the spatial and phenotypic information of every cell in the mIF image. Second, a unsupervised deep learning algorithm is applied, using graph convolutional networks (GCN) [10], to generate embeddings representing each cellular neighbourhood. Third, the embeddings are non-linearly down-projected into two dimensions, via uniform manifold approximation and projection (UMAP) [12], in such a way as to retain as much of the interesting structure in the embeddings as possible. Finally, the down-projected embeddings may be visually examined or statistically analysed for clustering and structure; noting that closeness in embedding space represents similarity between cellular neighborhoods in the highly complex, but real-world, spatio-phenotypic-morphological space of the mIF image.

In other words, GraphITE is a graph-based AI pipeline for converting mIF images into plots, statistics, and diagrams that a pathologist can quickly examine to identify hidden patterns in the data. This allows pathologists to gain new biological insights, deepening their understanding of the TME, and leading to a faster turnaround and better decision making in both drug development, and patient-therapy matching.

## 2 RELATED WORK

Graphs are mathematical structures that model pairwise relationships between entities. The entities are represented as nodes in the graph and depending on whether they interact with another node, an edge can be formed between them. Graphs can be used to model many types of interactions, processes, and relations in physical, biological and information systems. For example, in protein-to-protein interactions [21]. Graphs are also commonly in social media to model users’ relationships and to recommend content. Graphs are also found in geographical settings such as in digital maps, where the nodes might correspond to addresses or locations, while the edges can represent the distances between them. The use of graphs in this way can allow one to find shortest route to a particular destination of interest [3].

The use of graphs as a means to derive insights in the biomedical domain is not new, in fact, there are several successful attempts at leveraging the technology. In [20] colo-rectal cancer grading is carried out by transforming histology images into a graph, in which cell nuclei are represented as nodes in the graphs and links made based on node similarity. Training on these with a GCN they are able to reach state of the art grading accuracy.

Another example may be found in [6] where an attempt is made to circumvent the issue that standard CNNs do not account for all of the intricate features and spatial arrangement of cells in histology images. Building classifiers for detecting cancer can thus become more difficult. Their approach to the problem is to instead convert the H&E stained image into a graph (with cell nuclei as nodes and edges based on distances). Using graph convolutional neural networks in a supervised manner they train a model that reaches comparable performance with other, larger CNNs.

In another example [11] a GCN is employed in order to perform prediction of the status of human epidermal growth factors; H&E stained slides of breast cancer were transformed into a graph; and the approach was demonstrated to be both more computationally efficient and more performant than prior state of the art.

In [5] the DeePaN framework is described, where unsupervised learning is carried out on graphs integrating genomics and electronic health records data together, in order to identify responsive and non-responsive patient subsets amongst IO therapies addressing Non Small Cell Lung Cancer (NSCLC).

## 3 METHOD

This section describes the steps that constitute the graph-based pipeline, including data preparation, graph construction, graph embedding training, down-projection, and analysis.

### 3.1 Data Preparation

To demonstrate this pipeline, we used a dataset of 65 commercially-obtained samples of mIF stained tissue from (NSCLC) patients. The biomarkers used to stain this mIF tissue were DAPI, CD8, PDL1, CD68, PD1, Ki67, and CK [15]. Slides were imaged with a Vectra Polaris multispectral imaging platform (Akoya Biosciences ®) using OPAL fluorophores 520, 540, 570, 620, 650, and 690. The autofluorescence channel was also acquired and retained throughout the image analysis. Image analysis was conducted using HALO software Highplex FL module (Indica Labs ®). Whole slide results were exported with object data inclusive of positivity status (0/1) for each marker and location for each cell identified. Additional morphological features were also extracted, in the form of for example cell and nucleus area. These data were combined in a tabular format listing, for each cell ID, the X & Y spatial coordinates, the full set of biomarker immunofluorescence intensities, and the morphological features. From this data, additional morphological features, such as cell eccentricity could also be computed and added.

Due to the fact that not all of the samples had been created at the same time, under the same staining procedure and image capturing settings, the intensities of the detected dyes had to be normalized on a per batch level to ensure a common dynamic range. A standard mean-variance normalization was applied to each dye individually batch-wise.

In addition to the raw intensities of the biomarkers, the HALO image analysis system also reported whether a cell was deemed positive for a particular marker or not, based on built-in domain knowledge and certain rulesets. This marker positivity was encoded in the form of a boolean value for each of the markers along with the intensity. In order to try to capture both of the two values, the intensity and the boolean, a combined approach was taken as a preprocessing step in which the intensity values were normalized to fall in the range [0.0, 0.5] with an additional 0.5 added if the marker was positive. This ensured that the dynamic range of the intensity was preserved while at the same time making use of the built in domain knowledge in the cell detection tool. An example of the resulting data is shown in table 1. In total, about 45 million cells were processed.

**Table 1:** Example mIF data table including normalised biomarker intensities, and auxiliary features.

| Cell_ID | PDL1 | CD8 | Ki67 | CD68 | CK | PD1 | auto | DAPI | eccentricity | cell_area |
| --- | --- | --- | --- | --- | --- | --- | --- | --- | --- | --- |
| 265205 | 0.023221 | 0.077006 | 0.024235 | 0.062387 | 0.000517 | 0.009603 | 0.045871 | 0.504963 | 1.307692 | 171.745 |
| 297948 | 0.018611 | 0.074583 | 0.012308 | 0.023856 | 0.000480 | 0.019373 | 0.044164 | 0.528180 | 1.076923 | 143.162 |
| 313582 | 0.013624 | 0.043023 | 0.008150 | 0.045226 | 0.000580 | 0.009647 | 0.027494 | 0.504590 | 1.071429 | 168.049 |
| 117385 | 0.028539 | 0.038532 | 0.006990 | 0.082494 | 0.000196 | 0.031632 | 0.044176 | 0.506989 | 1.090909 | 100.287 |
| 387327 | 0.022115 | 0.032659 | 0.011969 | 0.086545 | 0.000484 | 0.007019 | 0.027095 | 0.504594 | 1.083333 | 127.638 |

### 3.2 Graph Construction

Since we are concerned with the interactions between cells in the TME, we construct graphs by mapping each cell in the mIF image as a node in a graph, using the X, Y coordinate as reported by the image analysis tool and connecting nodes with edges according to their distances. The selection of edges based on cell to cell distance inherently assumes a range over which cells can interact, either by direct contact or paracrine signaling with soluble factors. From a biological perspective, it is reasonable to focus on local interactions because these interactions in the TME, have been shown to influence not only the immune response to tumor through antigen presentation but also the active suppression of the adaptive immune response through expression of checkpoint inhibitors by tumor and immune cells [14][16]. However, it is worth considering carefully the distance thresholds at this stage since they constitute the first significant hyperparameter in the pipeline, and one which likely has a substantial impact on the embeddings generated and the insights derived thereby. Optimization of these thresholds may indeed be different for exploratory as opposed to predictive applications. An example of a mIF image with constructed graph overlay is shown in Figure 1.

**Figure 1:**
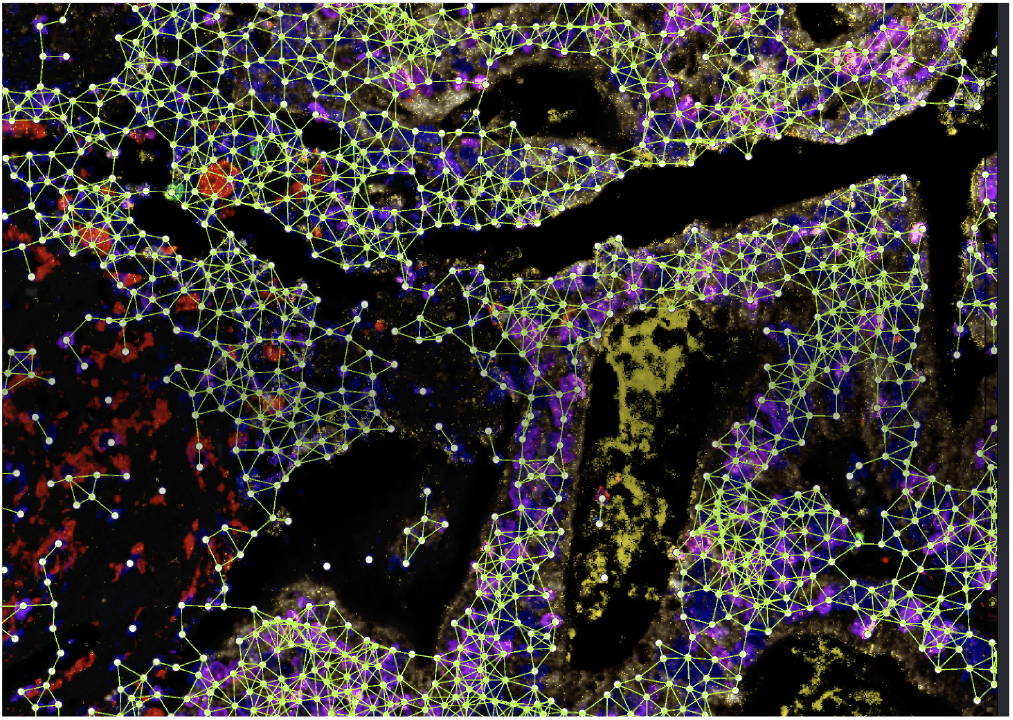
A mIF image with a constructed graph shown as an overlay.

As for the node feature values, we assign all (or a sub-set) or the available phenotypical (i.e. normalised biomarker immunofluorescence intensities) and morphological (i.e. cell eccentriciy and volume) features. The edges that connect nodes get assigned a weight inversely proportional to the distance between the two cells as 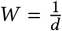

### 3.3 Deep Unsupervised Learning on the Graph

There are several ways in which embeddings can be generated from graph-based data. It can either be on the graph level, where one tries to represent the entire graph as an embedding, or at a sub-region or node level. The former can be useful when comparing different types of graphs or when classifying them in a fixed set, such as in [4]. The latter is useful when a more fine-grained area of the image is to be represented and explored, which is our purpose. One could imagine that given a dataset where response is known on a per tissue basis, that the full graph embedding could be interesting from a prediction standpoint.

At this stage we have one graph per mIF image, and it becomes possible to apply any of a range of unsupervised learning algorithms set to operate on graph structured data. There are several promising candidates for capturing information in graphs, such as GraphSage [8], Deep Graph InfoMax (DGI) [18], Node2Vec [7], and Variational Graph Autoencoder (GAE) [9].

While a full and exhaustive comparison of approaches would be illuminating, a comparison of graph-based unsupervised learning methods is not the primary focus of this work. It is, however, worth taking a moment to consider the underlying intuitions around these comparative methods.

DGI, for instance, is a contrastive method; i.e. it aims to maximise the distinctiveness between pairs of graphs: the true graph and a corrupted alternative of the graph. This is achieved by setting up a classification problem, whereby a neural network (called the discriminator) aims to predict whether a presented embedding belongs to the true graph or a corrupted graph. If the discriminator is effective at identifying embeddings that belong to true and corrupted graphs, we know that the embeddings must contain the necessary information required to distinguish between these graphs. In other words, DGI, aims to generate embeddings that *capture the distinctiveness of the graph & cellular neighbourhood*, as compared with corrupted alternatives. Naturally, this entails that careful thought be given to the choice of the so-called corruption function and readout function; since this choice changes what it means to say that a graph’s embeddings are *distinct*.

GraphSage uses a very similar approach in the way embeddings are generated for each node, but differs in the learning objective. In this case, the idea is to ensure that nearby nodes in the graph are given similar embeddings while distant nodes get very distinct from each other. This then puts emphasis on the assumption that nearby nodes are similar and should cluster together. While this is a sensible assumption from one perspective, it can mean that repeating patterns of nodes that are far away from each other, but otherwise similar, could end up with very different embeddings.

Conversely, GAEs work in fundamentally the same manner as all autoencoders: the goal is to output a prediction of the input via a bottleneck which hopefully captures a good representation of the latent space. In other words, an effective GAE aims to *capture and represent compactly the information necessary to reconstruct the graph*. This has the benefit of effectively guaranteeing no loss of information (under ideal conditions). Other methods, such as contrastrive methods, do not necessarily make this guarantee; however the loss of information may be measured via a proxy task we describe later in the experiments section.

Node2Vec builds upon Word2Vec, effectively treating random walks in the graph like words in a vocabulary. Word2Vec, and specifically skipgram negative sampling, trains embeddings that represents the likelihood that another work will be nearby the target word. The underlying assumption is that nearby words should be semantically related; i.e. if words appear near each other, they are probably related at a conceptual level as well. Thus, the embeddings generated using Node2Vec capture information such that other embeddings will be nearby if the probability distribution of random walks from the target cells are similar. Thus, cells that are nearby in the graph will always be somewhat nearby in the embedding space because somewhat similar random walks are always possible; however cells that are distant in the graph may also have nearby embeddings if the cellular neighbourhood is similar.

In this work we have utilised the DGI algorithm to generate node embeddings. We expect that similar performance is achievable via the discussed alternatives; but we have yet to fully explore the relative performance impact. Given the use of a contrastive approach, we consider it worthwhile to additionally evaluate the loss of information retained by the embeddings via proxy tasks, which we discuss later in the results section.

Under DGI, we utilised a row-shuffling corruption function; effectively randomising the connections between nodes in the graph, and an average embedding readout function as suggested in the orginal paper. This means that the embeddings are trained to be easily distinguished from neighbourhoods that have the same features (i.e. phenotypic or morphological information) but are connected in a different way.

An important hyperparameter in the embedding generation is the selection of GCN depth. With GCN layers, as with all convolutional layers, increasing depth translates directly to increasing receptive field. A one layer GCN integrates information from every node connected by an edge to the target node, whereas a two layer GCN additionally incorporates the information that updated those neighbouring nodes (i.e. their neighbours), and so on. In other words, each embedding incorporates information about nodes that are *N* hops way, where *N* is the number of layers in the GCN. The choice of GCN depth in the embedding generation of a tissue graph therefore dictates the effective ‘size’ of the cellular neighbourhood that each embedding represents; even though the embedding is always ‘focused’ on the target cell. The relative importance of the information in the target cell, compared with the information in closely, and more distantly, neighbouring cells is dictated by the interaction between several hyperparameters, namely: The distance threshold in graph construction, the edge weighting (if any), and the GCN depth. Together these hyperparameters dictate the meaning of the embeddings, which necessarily sit on a smooth scale ranging from purely cell embeddings, at the ‘narrow extreme’, to being driven by the average statistics of the entire graph, at the ‘broad extreme’.

The cellular neighbourhood size may be usefully guided by the pathologist or end-user; however in general there is a range that seems broadly applicable as most cells of interest in the tumor microenvironment range from 7-20 um in diameter for lymphocytes to 15-50 um in diameter for tumor cells [1] [13], and make up the TME alone with blood vessels, fibroblasts, and components of the extracellular matrix [19]. Specifically, since both extremes seem to miss a large part of the value of the method, (the ‘narrow extreme’ ignores the cell-cell interactions in the TME, whereas the ‘broad extreme’ approximates simpler graph-level statistical approaches), we aim for a set of hyperparameters with an effective radius of 30-300 microns for a cellular neighbourhood.

We selected an embedding size of 64 for our pipeline. Experimenting with this parameter showed that 64 represents a reasonable trade-off between loss of information and efficient computability.

The output of this stage of the pipeline is one embedding per cell in every mIF image. Importantly, via the graph convolutions, an embedding should not be interpreted as an embedding for a single cell, but rather an embedding of a cellular neighbourhood focused on a target cell.

### 3.4 Embedding Down Projection

With well trained 64-dimensional cellular embeddings, we can now conduct analyses to explore previously hidden patterns in the data. While there are useful analytic techniques that can be applied directly, such as clustering algorithms in 64-dimensional space, it is easier to understand the broad structure of the data by down-projecting and visually inspecting the embeddings. This is especially true when taking a hypothesis-free approach to data analysis.

Prior to visualisation of the trained embeddings it is necessary to project down to a lower dimensionality ideally two dimensions. Many algorithms exist for down projection, but linear methods (such as principal component analysis, or singular value decomposition) can often fail to capture complex manifolds in high dimensional space which may represent a pattern in the data that we aim to expose. More complex non-linear approaches such as t-distributed stochastic neighbour embeddings (TSNE) [17], and uniform manifold projection and approximation [12] can in such cases be more appropriate. For our pipeline we utilise UMAP, although we expect that other non-linear methods may perform similarly as well.

Because the total number of generated embeddings is very large (about 44 million), in order to make the UMAP computation feasible from a time-complexity perspective, a down-sampling is performed in which only a subset of 1000 random cells (their 64 dimensional embeddings) were selected per sample, and run through the UMAP algorithm.

### 3.5 Visualisation and Analysis

With the embeddings down-projected into 2D space, one can finally start to investigate whether any apparent clustering is visible. To this end we developed a tool for interactively exploring the UMAP while filtering by mIF image, generating subset biomarker statistics on-the-fly, and showing cellular neighbourhood inspection, as shown in fig 2.

**Figure 2:**
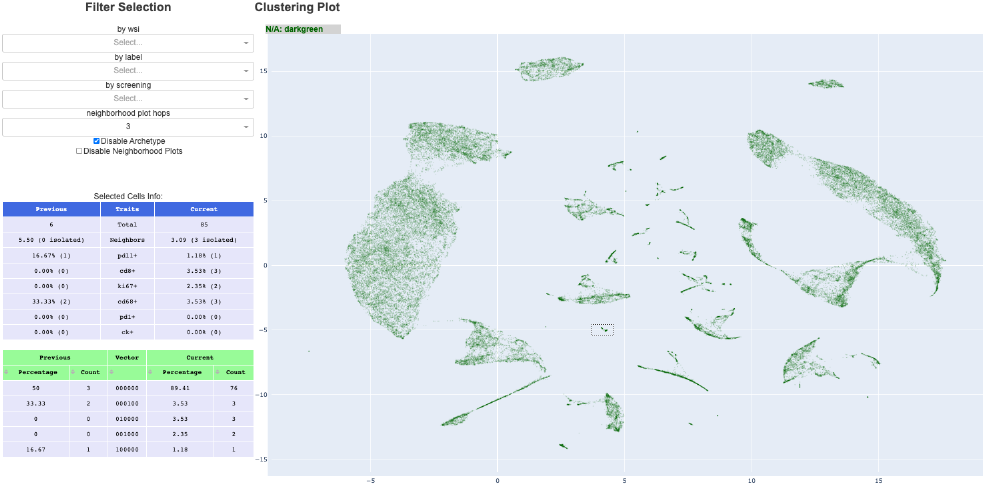
Embedding exploration tool showing down projected embeddings, summary statistics, selected embedding cellular neighbourhoods, and data filtering options (best viewed electronically).

For a demo, visit the following link where some of the functionality is showcased. https://az.box.com/v/mif-demo-vis

## 4 EXPERIMENTS AND RESULTS

This section details experiments carried out to validate the pipeline and perform initial exploration on our data to demonstrate how this tool could be used to investigate the biology contained in the mIF images.

### 4.1 Reconstruction experiment

With the embeddings generated, it is of interest to verify that we have not unduly lost information contained in the original mIF images. Unfortunately, many standard approaches to evaluating these aspects are not directly applicable to this problem. For example recall@K and MAP@R require labels for the embeddings, and it is not obvious what would constitute sensible labels for our dataset. Similarly, clustering approaches require specifying hyperparameters (such as number of clusters, or maximum distance) that are difficult to choose for arbitrary data. A measure such as trustworthiness is useful for estimating the loss of structured information that occurs during the down-projection stage, but does not help us to identify the loss of information captured via the embedding training itself.

To at least ensure that the information of the target cell of each embedding had been maintained, a simple experiment was devised aiming to demonstrate how well the initial information in the graph could be reconstructed from the trained embeddings. In principle, the embeddings should have captured this information, along with neighbouring cell information, so reconstructing the target cell features from the embeddings should be relatively simple. In particular in the case of the weighted graph, for each embedding, the emphasis would have been mainly put on the target cell.

To investigate whether the embedding still contained the original information, we train a single fully connected layer neural network to project from the embeddings to predicted target features (i.e. biomarkers). We train this layer on the training data, and report performance on the remaining validation dataset. These two sets of data were formed by simply splitting the full dataset in a 80-20 manner. Table 2 shows the performance statistics on this task, which demonstrate that the target node features could be reconstructed from the trained embeddings to a great degree. This verifies that indeed the target cell information is not lost as a result of the GCN step.

**Table 2:** Mean absolute error on the reconstruction task.

| DAPI | PDL1 | CD8 | Ki67 | CD68 | CK | PD1 | Auto |
| --- | --- | --- | --- | --- | --- | --- | --- |
| 0.0038 | 0.0047 | 0.0053 | 0.0064 | 0.0037 | 0.0029 | 0.0025 | 0.005 |

### 4.2 Effective Receptive Field

As discussed in the methods section, several hyperparameters play a crucial role in determining the receptive field (i.e. the size of the cellular neighbourhood) captured by the embeddings. The effective receptive field of the embeddings (in terms of microns on the tissue sample) is a non-trivial combination of three hyper-parameters: edge-connection distance threshold, edge-distance weighting, and GCN depth.

The edge-connection distance threshold has a profound effect, determining the very structure of the graph. The effect can be considered as a special case of edge-distance weighting however, where the weighting is a binary function of distance. The edge-distance weighting affects, more smoothly, the relative weighting of the more distant cells and the nearer cells in the neighbourhood. In effect, this modifies the kurtosis of the distribution of weighting across the the receptive field.

Finally, the GCN depth changes the number of hops that can affect the embedding at a target node; in effect ‘lifting’ the distribution of weighting upwards with more hops. Thus this parameter affects the total size of the receptive field without changing the shape of the distribution of weighting across it.

It is crucial to understand the interplay between these three hyper-parameters to develop a good intuition of the changing meaning of the trained embeddings. Figure 3 shows the interaction between these hyper-parameters in our experiments. In a future implementation of the downstream visualisation tool, it may be beneficial to map these interactions to a single parameter that the pathologist or end-user can control. Since the end-user likely does not need to grasp the nuances of these three hyper-parameters, but may very well care about the receptive field of the embeddings captured.

**Figure 3:**
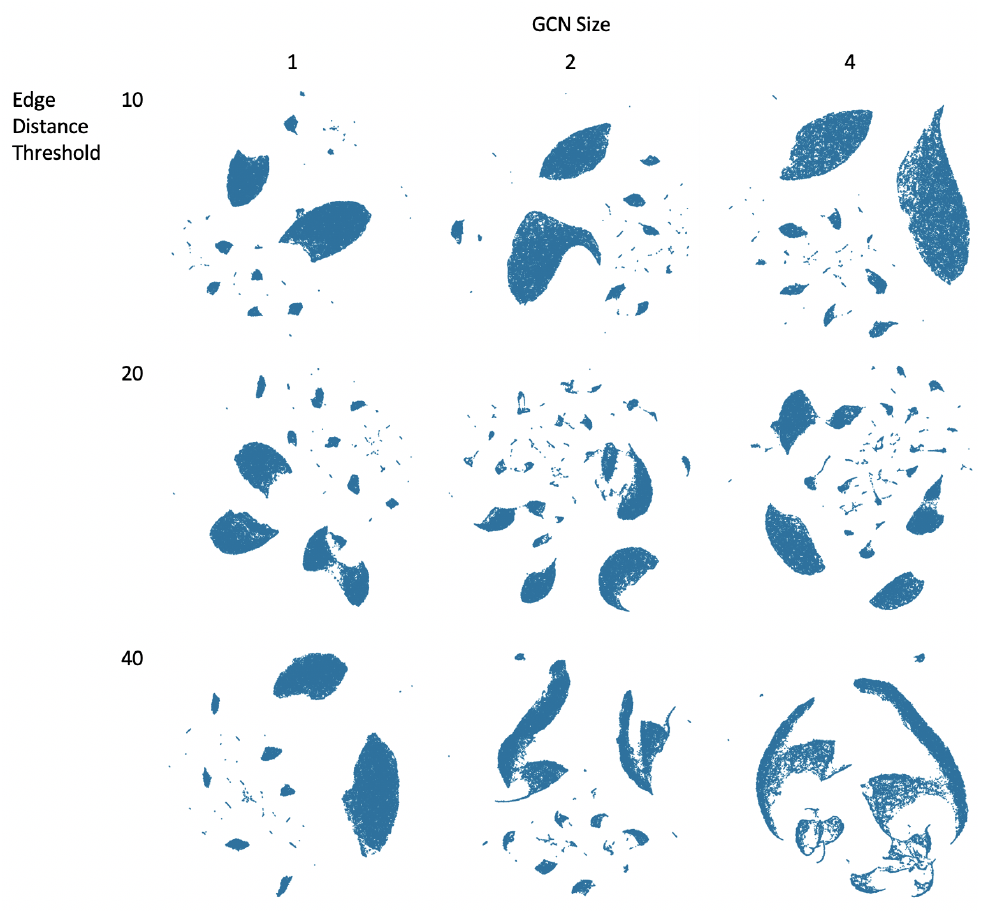
Down-projected embeddings for a set of different receptive-field-related hyper parameters (best viewed zoomed in).

**Figure 4:**
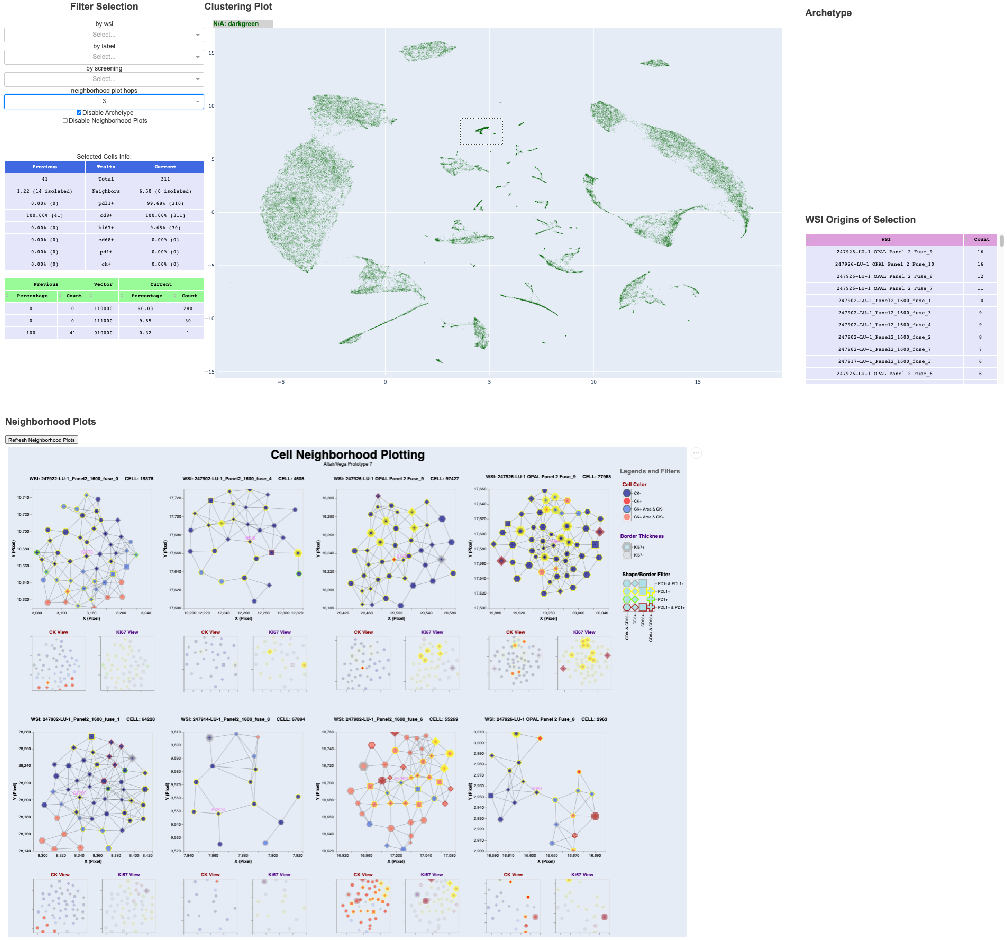
Initial exploration of the down projected embeddings shows several large clusters, as well as smaller ones. Each one seems distinct in terms of density of neighborhoods or available markers for the target cell. One cluster of cells is selected, and its distribution and a selection of neighborhoods are visible (best viewed electronically).

From the figure, it is clear that the majority of the clustering remains relatively unchanged for certain selections of hyperparameters. Indeed, as already mentioned, certain combinations of settings might actually approximate one another. For example the interplay between hops and distance thresholds. The fact that the clustering is relatively robust to these changes is quite welcome though, as it indicates that the particular choice made is not massively important for the outcome.

### 4.3 Exploring the data for biological meaning

Here we explore the down projected embeddings to demonstrate an example walkthrough of how to use this pipeline to explore the biology from a new perspective.

The place to start with embedding exploration is to view the down-projected embeddings and seek structure in the form of clustered groups of embeddings. In 4 we show a UMAP of embeddings trained on our dataset.

One can immediately observe several islands where cell embeddings seems to cluster together. Some larger, some smaller. Via the tool once can manually start circling the areas in order to get a feeling for what cells reside there. Having looked around for a while, the map shown in Figure 5 could be constructed. Two of the main things one may observe are the two major islands on either side of the plot. The one on the left hand side seems at first glace to be quite odd, with all of the markers negative. Although on second thought this is quite interesting, because that just means that there are several cells in the tissue that the mIF dyes simply do not characterize! A similar arguments can be made for the island on the right where the only difference is that the all the target cells are tumor (CK+). Given a set of additional markers, one would likely have observed clustering effects also within these major islands.

**Figure 5:**
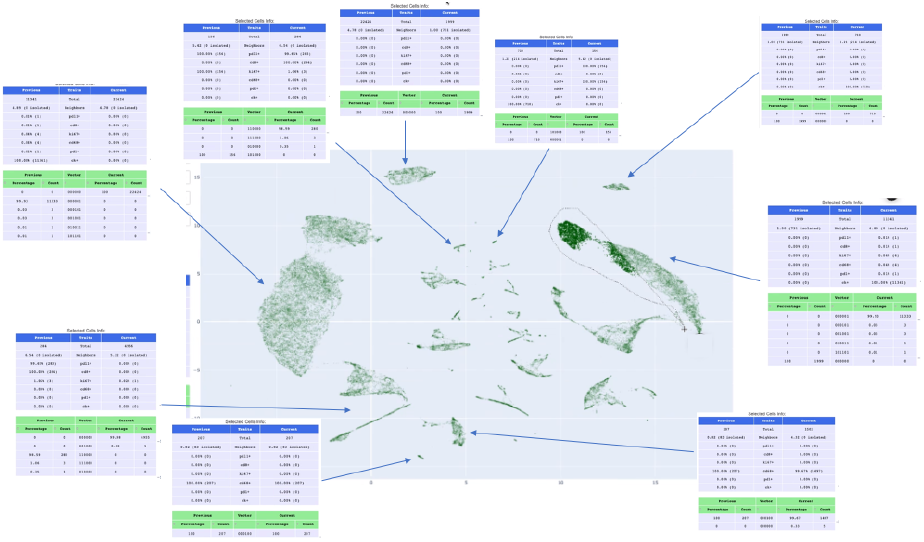
Summary statistics regarding some of the visible clusters in the UMAP visualizer. Part of an active lasso selection is visible to the right (best viewed electronically).

Continuing to investigate the central part of the UMAP reveals several clusters where a set of two or three other biomarkers are active. Interestingly enough, one can occasionally observe unexpected combinations of markers “in the wild”. While observations of rare multimarker phenotypes have been documented [2], the current accuracy of cell segmentation in image analysis software requires that unexpected phenotypes be visually confirmed on the image, preferably by a pathologist or other subject matter expert. This tool allows for the enumeration of many individual cells along with their coordinates to locate them within the original tissue sample. This has utility both for QC and for the identification of rare and potentially predictive or prognostic cell phenotypes or spatial phenotypes (small networks of cells).

Further exploration seems to continue to yield clusters where the major distinguishing factor is two or three biomarkers. A question naturally arises as a result from this. Does the system simply group cell embeddings based on their target cell characteristic? Luckily enough, this does not seem to be the case. Looking around the UMAP some more, one can observe that some clusters actually share common target cell features, but differ on things such as neighborhood density. Or, by actually using the cell neighborhood visualization feature of the exploration tool, one can see that indeed the neighborhoods in which the target cells finds themselves in are different.

A factor that seems to play an important role in the clustering of the cell embeddings is the neighborhood density. Notable for some islands is that the density is quite low, sometimes even having a good amount of cell neighborhoods just consisting of a single loner cell. It has been known for some time that density might play a role in how the tissue can behave in response to some drugs, but to carefully, not to say consistently, measure the density between samples is very demanding for a human pathologist. Having a way to characterize the cell density in an automated way like this, and to be able to explore such neighborhoods might be helpful.

All in all, the tool in its nascent form does not give direct answers or point in any particular direction. It does, however provide the user with a new perspective of the data. These new ways of visualizing the TME and investigating the clusters of cell neighborhoods are an incredibly valuable tool for understanding how therapies, time, or demographics may affect consequential cell populations in and around tumors, as well as in other contexts in which more complete understanding of the interaction of multimarker phenotypes and cellular spatial arrangement may fuel the generation of novel testable hypotheses. Conducting experiments to verify these new hypotheses might in turn yield new data that can be visualized in a circle of incremental refinement of ideas.

## 5 CONCLUSION

The aim of this study is to explore the feasibility of an unsupervised graph-based network for generating embeddings of cellular neighbourhoods. The generated embeddings must be compact representations of complex high-dimensional mIF data that can be easily explored to provide new perspectives of the data, or test existing hypotheses.

We have shown that our pipeline can produce embeddings which retain the majority of the information in the original data and are robust to a range of sensible hyperparameter settings. The down projected embeddings were explored via a hypothesis-free or hypothesis-generating procedure, and this demonstrates how the pipeline offers a valuable approach in getting a new perspective on the TME as captured via the complex high-dimensional mIF imaging data.

## ACKNOWLEDGMENTS

We would like to thank our colleagues in the AstraZeneca mIF team for their valuable feeback, comments, and assistance with data provisioning. In particular Helen Angell, Lorenz Rognoni, Andreas Sptizmüller, and Günter Schmidt.

